# Mortality in an alpine ungulate increases nonlinearly with snow depth

**DOI:** 10.64898/2026.09.21.753322

**Authors:** Kallan Crémel, Marco Festa-Bianchet, Alexandre Langlois, Fanie Pelletier

## Abstract

Heavy snowpack can be a severe constraint for alpine herbivores, yet adult survival in long-lived temperate ungulates is typically canalized against environmental variation. Using a 45-year individual-based dataset on bighorn sheep (*Ovis canadensis*) and historical snow reconstructions at Ram Mountain, Canada, we evaluated the direct and indirect effects of snow conditions on age- and sex-specific survival. For both sexes, survival of lambs and senescent individuals decreased with increasing average snow depth, whereas non-senescent adults (aged ≥ 1 year and < 8 years) showed high survival under mean conditions. However, this resilience broke down once specific thresholds of snow depth were crossed, with sharp survival declines beyond 90 cm of snow depth for females and 100 cm for males. Lamb survival, already reduced by increasing average snow depth, declined further once snow depth exceeded 90 cm. Crucially, the annual probability of snow depth exceeding 90 cm increased markedly over the study period, along with a density-independent temporal decline in survival across all demographic groups. Lamb survival declined by 0.37, and adult female survival, the most critical vital rate for population growth, dropped by 0.13 over the study period. Our findings challenge the established paradigm of demographic buffering in prime-aged adults, demonstrating that their resilience to environmental variability is fundamentally non-linear and breaks down once a changing climate regime breaches adult physiological boundaries, severely compromising population viability.

## Introduction

Mountain and high-latitude ecosystems are characterized by extreme seasonality and prolonged snow cover. For long-lived species inhabiting these environments, including ungulates, deep snow can be a severe physiological and mechanical constraint that shapes population dynamics (Horne et al., 2019; Jackson et al., 2021; LaSharr et al., 2023). Winter environments impose energetic constraints on individuals through the high metabolic demands of thermoregulation in cold temperatures (Parker & Robbins, 1984). Deep snow compounds this by increasing the metabolic costs of locomotion, especially when depth exceeds chest height (Dailey & Hobbs, 1989; Parker et al., 1984). When snow buries vegetation, it required individuals to invest energy in cratering through the snowpack to access low-quality winter forage (Fancy & White, 1985; Goodson et al., 1991; Parker et al., 2009). These nutritional and physical limitations are also compounded by an elevated risk of predation, as deep snow impedes ungulate escape efficiency while often increasing hunting success and travel efficiency in large carnivores (Huggard, 1993; J drzejewski et al., 2002; Sullender et al., 2023). Moreover, snow conditions can affect individuals through complex pathways that involve both direct and indirect effects (Crémel, Festa-Bianchet, et al., 2026; Hurley et al., 2014; Post & Stenseth, 1999). While snow acts as an immediate mechanical barrier to locomotion and foraging, it can also induce delayed seasonal carry-over effects by persistently altering body condition into the following autumn (Crémel, Festa-Bianchet, et al., 2026). Taking all these factors into account, one can expect that deep snow or changes in snow precipitation could abruptly affect ungulate populations, leading to sudden demographic shifts (Desforges et al., 2021; White et al., 2025).

To accurately capture the impact of deep snow events, the choice of metrics is important. Traditionally, studies rely on average environmental conditions (van de Pol et al., 2016) to assess effects on survival. While mean environmental conditions can assess general winter severity, they can mask possible non-linear constraints of extreme events. For example, the metabolic cost of locomotion in snow increases non-linearly and abruptly when snow reaches chest height (Dailey & Hobbs, 1989). A consideration of mean snow depth would dilute this biological signal, by failing to capture specific events, like the number of days spent in deep snow, which represent an important constraint for locomotion. The masking effect of climate averages is a recognized challenge in population ecology; for example, mortality in Soay sheep (*Ovis aries*) is driven by the occurrence of winter storms rather than average local weather conditions (Coulson et al., 2001). Ultimately, for populations facing variations in average winter severity or deep snow events, demographic responses are fundamentally driven by age-specific vulnerabilities.

In long-lived vertebrates, a decrease in juvenile survival is often a sensitive early warning signal of environmental stress (Eberhardt, 2002; Gaillard et al., 2000). Because young individuals face strict morphological constraints, they tend to allocate energy toward attaining a sufficient body mass for winter survival rather than in reserve accumulation (Parker et al., 2009). They are therefore more affected by resource scarcity. Adult survival, in contrast, is predicted by life-history theory to be stable, or canalized, against environmental stochasticity (Gaillard et al., 1998). Recent frameworks have formalized this buffering mechanism, yet identify nonlinear responses of demographic rates to environmental drivers as a critical, still underexplored, component (Gascoigne et al., 2025; Le Coeur et al., 2022). Given that adult survival contributes disproportionately to population growth rate, when conditions deteriorate, selection is expected to favor individuals that prioritize their own survival, by drawing on fat and protein reserves accumulated prior to winter (Monteith et al., 2013), over allocation to reproduction. Whether the buffering of survival over environmental challenges holds under increasingly frequent or severe events, however, remains poorly tested.

While the effects of winter on ungulate population dynamics are increasingly well documented (Portier et al., 1998; Post & Stenseth, 1999), most studies rely on temperature or precipitation as proxies, and comparatively few have explicitly examined snow conditions themselves as a distinct driver of survival. Fewer still have explored the possible effects of average snow conditions, snow depth thresholds, and carry-over effects linking weather in one winter to body condition in the next. This gap is particularly critical in the context of climate change, which is expected to generate novel winter conditions, limiting the ability of existing frameworks to forecast demographic responses.

Here, we overcome these limitations by integrating the reconstruction of historical snow records from the SNOWPACK model (Crémel, Langlois, et al., 2026) with long-term monitoring of individual bighorn sheep (*Ovis canadensis*) on Ram Mountain, Canada (Festa-Bianchet et al., 2019). This framework provides an excellent opportunity to evaluate how snow conditions affect survival of different sex-age classes. To do so, we first evaluated how snow cover duration, depth, and density influence survival across all demographic groups. Accounting for age and autumn body mass, we predicted that persistent, deep, and dense snow cover would reduce survival, particularly lambs. Second, we tested for critical depth thresholds that may lead to a nonlinear drop in survival. By comparing the number of days exceeding thresholds in snow depth ranging from 10 to 100 cm, we searched for a depth at which snow suddenly becomes a limiting factor for survival. Finally, building on previous findings that winter snow conditions can induce carry-over effects on body mass the subsequent autumn (see Crémel, Festa-Bianchet, et al., 2026), we wanted to disentangle the direct and indirect pathways through which snow conditions affect age- and sex-specific survival.

## Methods

### Study area and population monitoring

We study bighorn sheep at Ram Mountain in Alberta, Canada (52°20’N; 115°46’W). Located approximately 30 km east of the main Canadian Rocky Mountain range, the study area spans 38 km² of alpine tundra and subalpine meadows, at elevations ranging from 1430 to 2160 m (Cloutier et al., 2024; Jorgenson et al., 1993). This population is geographically isolated, resulting in negligible immigration or emigration, which has facilitated demographic tracking since 1971 (Festa-Bianchet et al., 2019).

### Data collection and animal handling

We captured sheep annually from May to September using salt-baited corral traps and weighed them with a Detecto spring scale (accuracy ± 250 g) (Larue et al., 2022). We standardized body mass to June 5 and September 15 using mixed-effects models, following Martin & Pelletier (2011). Survival was determined from annual censuses. Given that emigration is very rare (Jorgenson et al., 1997; Loison et al., 1999) and the probability of detection is very high (95% for males and 99% for females, Jorgenson et al., 1997), any individual not seen in one field season was considered to have died in the previous winter. Population density was the total count of ewes aged ≥ 2 years as in previous studies of this population (Portier et al., 1998). Since we reconstructed snow from 1979 to 2023, the final dataset used in analyses included those 45 years. All animal procedures were approved by the Université de Sherbrooke Animal Care Committee (Protocol FP: 2024-4569) in accordance with the guidelines of the Canadian Council on Animal Care.

### Snow characteristics reconstruction

We used a historical reconstruction of local snow conditions since 1979 using the SNOWPACK model. A comprehensive description of the methodology and validation is provided in Crémel, Langlois, et al. (2026). This model estimated several annual metrics, including snow cover duration, start and end dates of the snow season, and mean, median, and maximum values for snow depth (cm), snow density (kg/m³), and snow water equivalent (mm). However, *a priori* correlation analyses revealed multicollinearity among these variables (r > 0.6; Appendix S1: Figure S1). Consequently, we restricted our analyses to three weakly correlated metrics (between −0.20 and 0.46) that are biologically relevant to our hypotheses: snow cover duration, average snow depth, and median snow density. We also estimated the number of days when snow depth exceeded thresholds ranging from 10 cm to 100 cm in 10-cm increments.

### Statistical analyses

All modeling was implemented using R v4.5.1 (R Core Team, 2025). Continuous explanatory variables were z-scored prior to analysis to facilitate the interpretation of coefficients and aid algorithm stability. To account for sex-specific aging, a preliminary modeling step identified the optimal functional form for the sex-specific relationship between age and survival. For both sexes, this relationship was best captured using natural cubic splines (*df* = 3; Bates & Venables, 2022) (Appendix S1: Table S2). We developed separate model sets for three demographic group (lambs, females aged 1 year and older and males aged 1 year and older) to account for sexual differences in survival and the greater vulnerability of lambs (Gaillard et al., 1998). Within the two adult groups, we use the term ‘adult’ to refer to predictions adjusted to the mean age of individuals in our dataset (5 years for females and 3 years for males), whereas ‘prime-aged’, ‘yearling’, and ‘senescent’ refer to marginal predictions for the discrete age classes described below. This stratified approach simplified the models by avoiding high-order (three-ways) interactions between continuous variables. To capture the sex-specific survival trajectories of lambs and the expected higher vulnerability of males (Festa-Bianchet et al., 1996; Leblanc et al., 2001), the lamb models included interaction terms between sex and mass, as well as between sex and all environmental covariates.

Bayesian inference was conducted using the *brms* package (v2.22; Bürkner, 2017) with default weakly informative priors. For each model, we ran four Markov chains with 10000 iterations, including a 4000-iteration warmup phase. Model convergence was confirmed by visual inspection of trace plots and R_hat values, which were all exactly 1.0. In line with Vehtari et al. (2021), we ensured that bulk and tail effective sample sizes (ESS) exceeded 400. Model adequacy was validated through posterior predictive checks. Given the binary nature of survival, we performed posterior predictive checks using bar plots comparing observed and predicted frequencies. These checks confirmed that the models accurately reproduced the observed number of survival and mortality events (*i.e.*, the frequency of 0 and 1), indicating that model structure adequately captured this aspect of the data.

To evaluate the effects of snow conditions on survival, we implemented Generalized Linear Mixed Models (GLMMs) with a Bernoulli distribution with a logit link. We included year as a random intercept to account for shared environmental stochasticity and inter-annual survival fluctuations (Kéry & Royle, 2016; McElreath, 2018), including discrete, well-documented events such as years of intense cougar predation (Cloutier et al., 2024). Year as random effect account for year-specific mortality spikes unrelated to snow conditions, allowing the fixed snow-related effects to be interpreted independently of those episodic sources of mortality. We did not include an individual random effect (ID) because in discrete-time survival analyses with binary outcomes, death occurs only once per individual and is therefore an absorbing event. Including an ID-specific intercept would lead to perfect separation, as the random effect would perfectly predict the terminal event, leading to mathematical non-identifiability and preventing model convergence (Gelman & Hill, 2006).

We generated a distinct set of predictions for each demographic group (Appendix S1: Table S1). We assessed the models relative performance using leave-one-out information criteria (LOOIC) based on Pareto smoothed importance-sampling (PSIS-LOO) (Vehtari et al., 2017). Model selection was based on the expected log predictive density (ELPD), where the model with the highest predictive performance was retained.

To verify that conclusions were not influenced by prior choices, we conducted a power-scaling sensitivity analysis using the *priorsense* package (v1.2; Kallioinen et al. 2023). By quantifying the cumulative Jensen-Shannon distance under perturbations of both priors and likelihoods, we confirmed that the posterior estimates were robust and primarily informed by the empirical data.

To quantify the certainty of the trends observed for all analyses, we reported the probability of direction (pd). This Bayesian metric represents the proportion of the posterior distribution that shares the same sign as the median estimate (Makowski, Ben-Shachar, & Lüdecke, 2019; Makowski, Ben-Shachar, Chen, et al., 2019). We categorized the evidence based on the pd value as follows: parameters with a pd < 80% were considered as having little to no directional support; 80% ≤ pd < 90% was classified as a moderate trend; 90% ≤ pd < 95% indicated strong support for a directional effect; and pd ≥ 95% was considered highly credible evidence.

To investigate potential snow depth threshold effect on survival, we used a series of GLMMs for each demographic group (lambs, females aged 1 year and older and males aged 1 year and older), testing a sequence of threshold variables, defined as the number of days each winter when snow depth exceeded X cm (Days > X). We evaluated thresholds from 10 to 100 cm with 10 cm increments, resulting in 10 candidate models per demographic group.

To isolate the specific effect of the snow depth threshold, each model included as covariates the natural cubic spline of age, fitted separately within the adult female and adult male groups, along with autumn body mass, population density, and the median snow density for that winter. To account for unmeasured inter-annual stochasticity, we included a random intercept for the year. As for previous analyses, all models were fitted using the brms package. We ran four chains with each run for 10000 iterations, including 4000 warmup iterations. To determine the most biologically relevant snow depth threshold for each demographic group, we evaluated the candidate models based on their predictive performance. We computed the LOOIC, models with higher ELPD values were considered to provide a better fit to the data.

To model long-term temporal changes in the frequency of snow depth threshold variables across the study period, we analyzed the annual number of days exceeding each selected threshold over 45 years using beta-binomial regression models. This approach was selected to account for the bounded nature of the response variable (constrained by the annual number of days with snow cover), while naturally accommodating the overdispersion and high proportion of zero-count years in the dataset. Year was included as a continuous covariate, with the first year of data (1979) as zero, ensuring that the model’s intercept represents the estimate at the start of the study. We ran four chains with 6000 iterations each (including 2000 warmup iterations). Model diagnostics were evaluated using posterior predictive checks. We verified that the simulated data accurately reproduced the empirical mean, standard deviation, empirical cumulative distribution functions, and the frequency of zero values observed in the raw data. To further explore temporal trends, posterior draws were extracted to derive the annual probability of experiencing at least one day meeting the threshold criteria, calculated as 1 – P(Y=0).

Following our finding of long-term temporal increases in snow threshold frequencies, we formulated a *post-hoc* hypothesis to evaluate a temporal decline in survival over the 45-year study period. To test this, we modeled individual survival using a GLMM specified as before with a Bernoulli distribution and a logit link function. Because annual population density varied over the study, we fitted a density-independent trend model by explicitly controlling for population density. Incorporating this covariate blocks the indirect effect of density on survival, thereby isolating the direct, residual temporal trend in survival. To prevent temporal pseudoreplication and ensure a conservative test of the temporal covariate, we implemented a dual temporal structure. The long-term trend was modeled as a continuous effect of year, with 1979 set to 0, and stochastic interannual fluctuations were accommodated with a random intercept for the year. Furthermore, to account for potential shifts in the age structure of the population and in autumn individual body condition over time, individual autumn body mass and age (modeled with the same natural cubic spline as in previous models) were included. For the lamb model, sex was included in interaction with other covariables. Model diagnostics were evaluated using posterior predictive checks.

To quantify the pathways through which snow characteristics influence sex- and age-specific survival to the following spring while accounting for previous autumn body mass, we performed a G-computation analysis (Robins, 1986). G-computation is a counterfactual simulation framework that standardizes predictions over the empirical distribution of the population to estimate true marginal effects. Unlike traditional conditional predictions, which often isolate a hypothetical average individual by fixing covariates at their mean values, G-computation utilizes the actual individuals from our dataset. This approach preserves the observed covariance structure and natural heterogeneity of the study population, ensuring that the resulting survival estimates represents genuine population-level effects.

The predictive models estimating the effects of snow characteristics on autumn body mass were derived from previous work (Crémel, Festa-Bianchet, et al., 2026). We embedded G-computation within a counterfactual mediation framework (Pearl, 2021) to decompose the total impact of snow characteristics. By constructing nested counterfactual scenarios, we mathematically decoupled the direct environmental pressure of snow from its indirect consequences mediated through body mass. This allowed us to simulate specific ecological states, such as deep snow conditions paired with baseline autumn body mass, that are biologically impossible to observe concurrently in nature but are essential for quantifying their respective influence (Daniels et al., 2023; Hernán & Robins, 2025).

For each sex, we conducted simulations of survival rates for three representative age classes, yearlings (1 year), prime-age individuals (3 years), and senescent individuals (9 years) (Festa-Bianchet et al., 2019). The analysis followed a sequential predictive chain driven by a specific snow scenario. For an individual in year *t*, the experienced winter conditions determined the predicted spring body mass in year *t+1*, which subsequently dictated the predicted autumn body mass at year *t+1*. To accurately reflect the chronological progression of the life cycle, time-varying covariates such as individual age and population density were dynamically updated within our simulation. This predicted autumn mass for year *t+1* was then used as the primary input to predict the probability of survival to the subsequent year, with individual ages and population densities changed to match the next life stage.

To explore the causal path, we compared three distinct counterfactual scenarios against a baseline of minimum observed snow depth, duration, or density. The total effect of snow was estimated by allowing the snow variables to vary simultaneously across both the body mass and the survival prediction steps, thereby capturing the comprehensive impact of winter conditions. The indirect effect, mediated through body mass, was isolated by predicting survival under baseline snow conditions while forcing the model to use the lower body mass values predicted under high snow conditions. Conversely, the direct effect was isolated by predicting survival rates under high snow conditions while holding the body mass inputs constant at the values predicted under the baseline snow scenario. To ensure our estimates accounted for both environmental and individual stochasticity, we performed these simulations using 1,000 draws from the MCMC posterior distributions for the demographic groups previously identified as vulnerable to snow-induced carry-over effects.

## Results

For all demographic groups, the model without interactions between population density or age was the best-supported model (Appendix S1: Table S4). Because age was modeled as a continuous covariate within the adult female and adult male groups rather than as discrete age classes, results below are reported for the mean age in both sexes; age-specific effects for yearling, prime-aged, and senescent individuals are instead presented separately in the path analysis (Figure 5). Lamb survival was lower for males compared to female (pd = 100%) and decreased for both sexes at high population density (Appendix S3: Table S1, pd = 100%) but the negative effect of population density was lower for male compared to female lambs (Appendix S3: Table S1, pd = 98%). Among the snow variables, average snow depth had a strong negative effect on lamb survival (Figure 1A, Appendix S3: Table S1, pd = 98%), followed by snow cover duration, which had a moderate negative effect (Figure 1B, pd = 89%). The negative impact of snow depth was more pronounced for females than for males (Figure 1A, Appendix S3: Table S1, pd = 89%). Conversely, male survival was more affected by snow density (Appendix S3: Table S1, pd = 88%) with no effect on females (Appendix S3: Table S1, pd = 57%).

**Figure 1:**
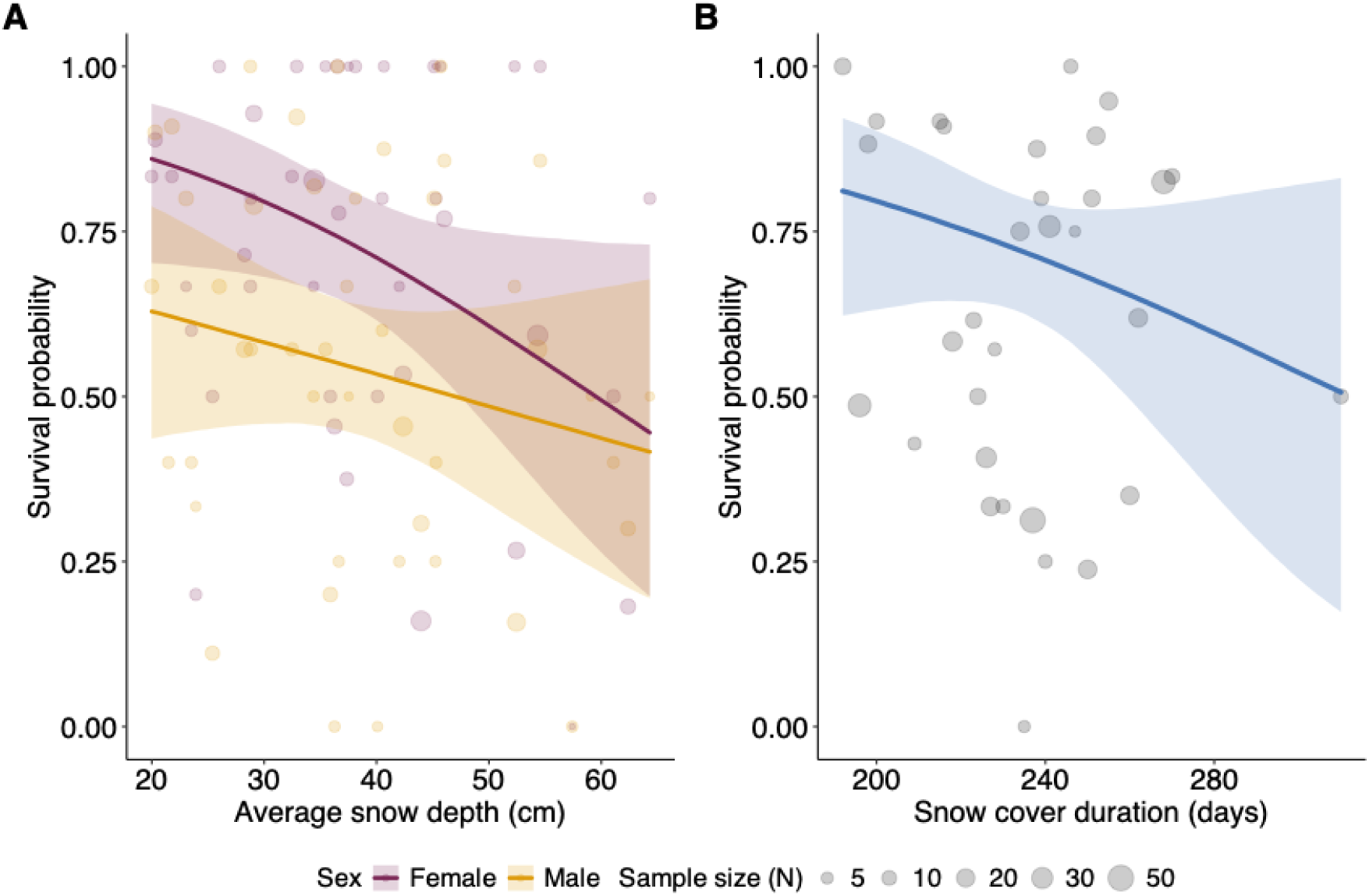
(A) Effect of average snow depth (cm) and (B) effect of snow cover duration (days) on lamb survival. Mauve line represents females and orange line males, with shaded areas indicating the 95% credible intervals, blue line represents the main effect fixed at reference level (female). Points represent the observed empirical survival proportions for each unique value of average snow depth and effect of snow cover duration, with point size proportional to sample size (N).

In contrast to lambs, adult female survival was largely unaffected by snow conditions and population density. While snow duration (Appendix S3: Table S1, pd = 51%) and snow density (Appendix S3: Table S1, pd = 63%) had no impact on adult female survival, average snow depth showed a weak negative trend (Appendix S3: Table S1, pd = 80%). Population density had no detectable influence on adult female survival (Appendix S3: Table S1, pd = 53%).

For adult male survival, average snow depth (Appendix S3: Table S1, pd = 84%) had a weak negative effect. Snow cover duration (Appendix S3: Table S1, pd = 75%) showed no clear effect, as well as snow density (Appendix S3: Table S1, pd = 63%) and population density (Appendix S3: Table S1, pd = 70%).

### Snow depth threshold

For lambs, a higher number of days with snow depth exceeding 90 cm severely reduced survival (β = −0.66, 95% CI = [−1.11, −0.22], pd = 100%, Figure 2A), though this negative impact was significantly attenuated for male lambs compared to females (β = 0.48, 95% CI = [0.05, 0.91], pd = 99%, Figure 2A). For adult females, the number of days with snow deeper than 90 cm also had a clear negative effect on survival (β = −0.38, 95% CI = [−0.60, −0.15], pd = 100%, Figure 2B) while adult males showed a stronger negative effect when snow depth exceeded 100 cm (β = −0.28, 95% CI = [−0.63, 0.05], pd = 96%, Figure 2C).

**Figure 2:**
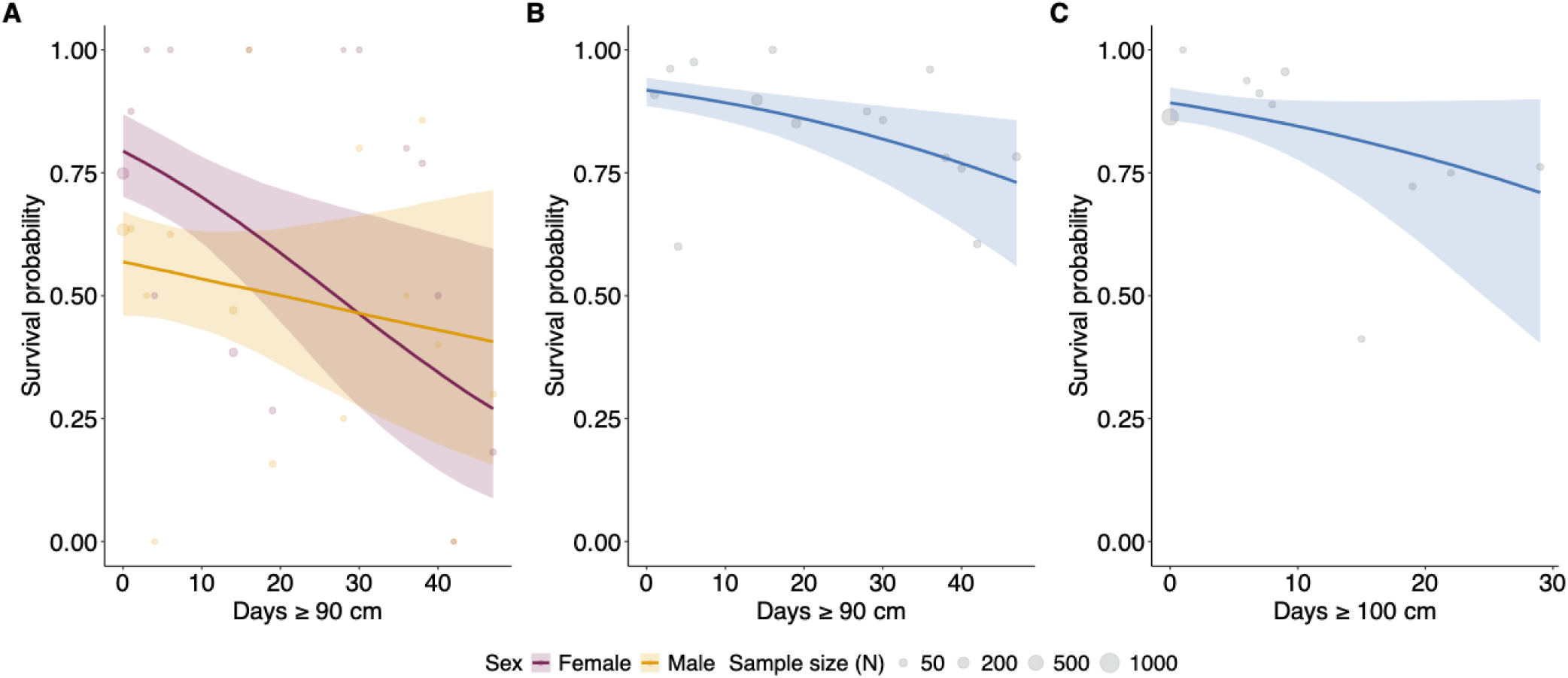
Effects of the number of days exceeding a threshold of snow depth on survival. Lamb (A) survival by sex (mauve line: female, orange line: male) relative to the number of days with snow depth ≥ 90 cm, and survival of adult females (fixed at mean age : 5 years old) (B) (≥ 90 cm) and of adult males (fixed at mean age : 3 years old) (C) (≥ 100 cm), respectively. The solid lines represent the fitted predictions, and the shaded areas indicate the 95% credible intervals. Points represent the observed empirical survival proportions for each unique number of days, with point size proportional to sample size (N).

Over 45 years, the annual number of days with snow depth exceeding 90 cm increased from 0 to 24 days, a strong positive temporal trend (Figure 3B, β = 0.08, 95% CI = [0.07, 0.08], pd = 100%). Concurrently, the probability of experiencing at least one day per year with snow depth above 90 cm go from 0.08 to 0.87 (Figure 3A). Similarly, the number of days with snow exceeding 100 cm increased from 0 to 9 (Figure 3D, β = 0.07, 95% CI = [0.06, 0.08], pd = 100%), while the annual probability of reaching this threshold increased from 0.04 to 0.61 (Figure 3C). Average snow depth and the number of days with snow depth exceeding 90 cm were strongly correlated (r = 0.71), indicating that winters with deeper average snowpack also tended to have more days above this threshold.

**Figure 3:**
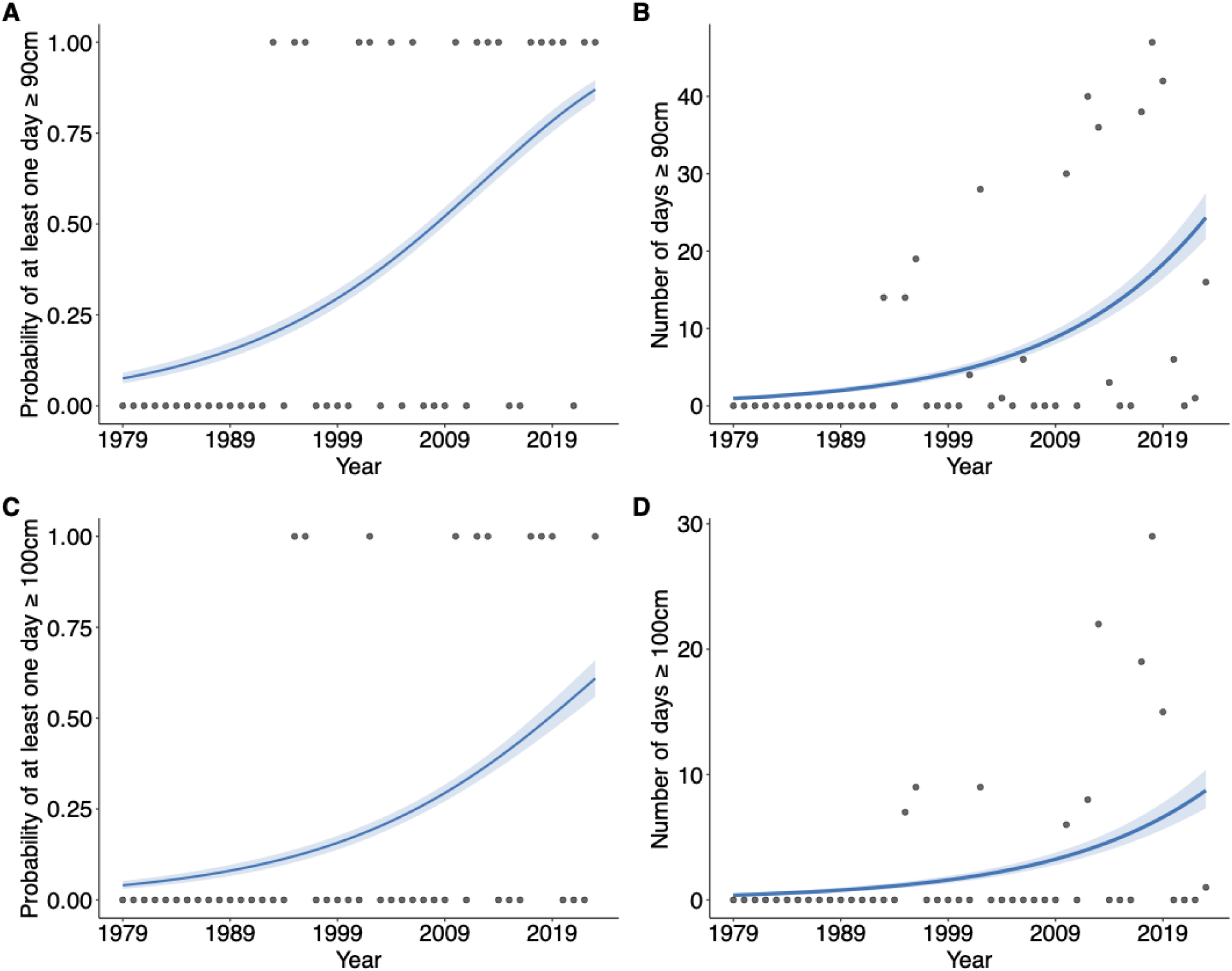
Temporal trends in deep snow conditions over 45 years (1979-2023), at Ram Mountain (Alberta, Canada). Panels show the annual probability of experiencing at least one day with snow depth exceeding (A) 90 cm and (C) 100 cm, alongside the total annual number of days with snow depth exceeding (B) 90 cm and (D) 100 cm. Blue lines represent the predicted mean trends from beta-binomial regression models, shaded areas indicate the 95% credible intervals, and points represent observed annual data.

### Temporal trend in survival

After accounting for population density, autumn body mass, and age, we observed a long-term temporal decline in survival over the 45-year study period, though the magnitude of this effect varied across demographic groups. The negative temporal trend was most pronounced in lambs, which experienced an estimated absolute decline in survival of 0.37 over the study period (Figure 4A, Appendix S3 : Table S4, pd = 98%). We found no substantial evidence that this temporal decline differed between male and female lambs (pd = 73%). Adult females also showed strong statistical support for a temporal decline (Figure 4B, Appendix S3 : Table S4, pd = 99%), corresponding to a 0.13 drop in survival. In contrast, adult males experienced a more moderate temporal decline, showing a 0.07 decrease in survival (Figure 4C, Appendix S3: Table S4, pd = 90%).

**Figure 4:**
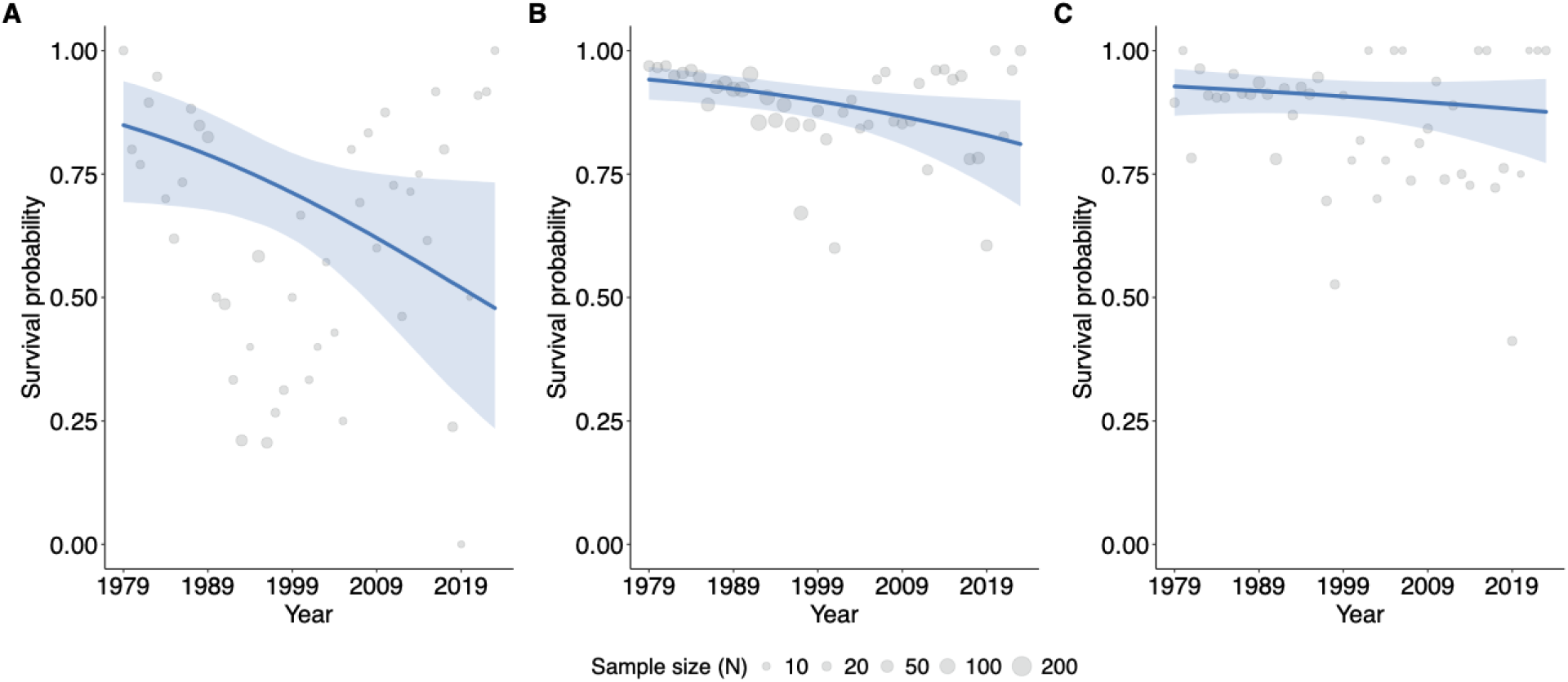
Effects of year on survival of bighorn sheep at Ram Mountain, 1979-2023. Panels show lambs (fixed at female level) (A), adult females (fixed at mean age : 5 years old) (B) and adult males (fixed at mean age : 3 years old) (C). The solid lines are fitted predictions, and the shaded areas indicate the 95% credible intervals. Points represent the observed empirical survival proportions for each year, with point size proportional to sample size (N).

### G-computation

The G-computation analyses revealed that average snow depth exerted the most consistent and pronounced negative marginal impact on survival, particularly affecting senescent individuals of both sexes (Figure 5, Appendix S3 : Table S5). For senescent males, the marginal total effect of snow depth was strongly negative (Figure 5, Appendix S3 : Table S5, pd = 86%), with a large portion of this impact being direct (Figure 5, Appendix S3 : Table S5, pd = 82%) and a smaller portion being indirect but less certain (Figure 5, Appendix S3 : Table S5, pd = 80%). Snow depth similarly affected senescent females, where the marginal total effect was also negative (Figure 5, Appendix S3 : Table S5, pd = 86%), driven primarily by its direct component (Figure 5, Appendix S3 : Table S5, pd = 82%). The marginal total effect of snow depth was also negative for yearling females (Figure 5, Appendix S3 : Table S5, pd = 92%) and yearling males (Figure 5, Appendix S3 : Table S5, pd = 84%). For prime-aged adults, the marginal total effect remained negative, with similar support for females (Figure 5, Appendix S3 : Table S5, pd = 87%) and males (Figure 5, Appendix S3 : Table S5, pd = 86%). Notably, we found a small marginal indirect effect of snow depth on female (Figure 5, Appendix S3 : Table S5, pd ranging from 86% to 99%), supporting the hypothesis of an indirect effect via autumn body mass. The indirect effect was less certain for male (pd = 80%). For males and senescent females, this indirect effect was less certain (Figure 5, Appendix S3 : Table S5, pd ranging from 80% to 86%).

**Figure 5:**
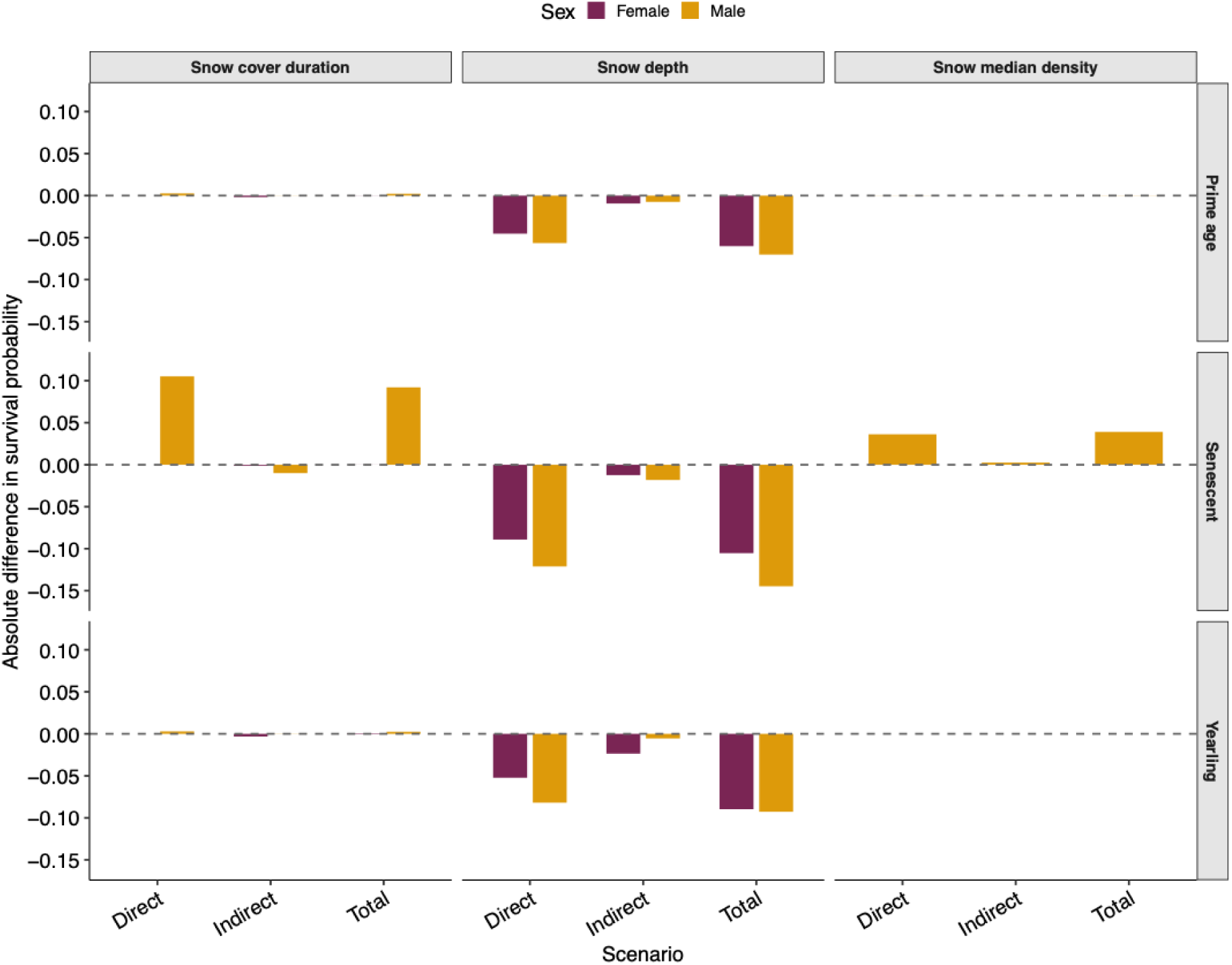
Absolute difference in survival of bighorn sheep associated with snow cover duration, snow median density and snow depth according to effect type (direct, indirect, and total), sex, and age class (yearling, prime age (3 years old), senescent (9 years old)) at Ram Mountain, Alberta, 1979-2023. Bars represent the absolute difference in estimated survival derived from G-computation, comparing a scenario where all individuals were exposed to the maximum versus the minimum observed value of each snow variable. The direct effect reflects the effect of the snow variable on survival independently of the mediator (autumn body mass); the indirect effect represents the portion of the effect transmitted through the autumn body mass; the total effect is the sum of both. Negative values indicate a reduction in survival associated with the snow variable. The dashed line marks the null effect (difference = 0).

In contrast to the effect of snow depth, snow cover duration and median snow density only had negligible marginal effects on survival (Figure 5, Appendix S3 : Table S5). For snow cover duration, the marginal direct and total effects for females hovered around zero with pd ranging from 51% to 69% (Figure 5, Appendix S3 : Table S5). For males, while the marginal total and direct effects of snow duration were positive, these estimates remained uncertain (Figure 5, Appendix S3 : Table S5, pd ranging from 75% to 78%). The largest effect for this variable was a negative but extremely weak marginal indirect effect in yearling females (Figure 5, Appendix S3: Table S5, pd = 99%). Finally, median snow density had no clear effect on survival (Figure 5, Appendix S3 : Table S5, pd ranging from 64% to 75% for prime-age and senescent males).

## Discussion

Our study revealed that the effect of snow conditions on alpine ungulate survival is not captured by average conditions alone. Rather, snow affects bighorn sheep survival through two distinct pathways: the overall severity of snow conditions and exposure to exceptionally high snow depths. While in years of high average snow depth the survival of vulnerable demographic groups such as lambs or senescent individuals of both sexes decreases, survival of prime-aged adults remains high. However, looking at critical snow depth thresholds reveals the limit of adult resilience. Over the 45-year study period, an increase in deep snow events has occurred in parallel with a density-independent temporal decline in survival across all demographic groups, including a major drop for adult females. Ultimately, this temporal decline reveals that adult survival canalization is not absolute but constrained by clear environmental boundaries when thresholds are crossed.

Susceptibility to average snow depth was more pronounced in lambs than in senescent individuals, a pattern that supports the high vulnerability of juveniles to external conditions in large herbivores (Gaillard et al., 1998). Lambs are also negatively affected by snow cover duration. Lambs have a smaller body size and shorter legs than adults, so they may experience higher energetic costs of locomotion in deep snow (Parker et al., 1984; Telfer & Kelsall, 1979). Furthermore, because lambs mostly allocate resources to growth, they enter winter with limited somatic reserves compared to adults. Thus, any restriction in dry matter intake or increase in daily energy expenditure compromises lamb growth (Crémel, Festa-Bianchet, et al., 2026) and ultimately survival. Interestingly, snow effects on lamb survival were sex-specific: average snow depth had a stronger negative effect on females, while males were more sensitive to snow density. This likely reflects morphological and energetic differences: smaller female lambs face a proportionally greater locomotor cost from snow depth (Festa-Bianchet et al., 1996), whereas dense, hardened snow more strongly restricts forage access for male lambs that likely have greater energy requirements that likely have greater energy requirements (Fancy & White, 1985; Leblanc et al., 2001). Notably, males showed lower average survival but a weaker decline under deep snow than females, potentially reflecting stronger pre-winter selection against frail males (Clutton-Brock, Albon, et al., 1985).

As capital breeders, adult bighorn sheep enter the winter with substantial somatic reserves (Festa-Bianchet et al., 1998; Jönsson, 1997), allowing them to buffer average snow conditions by drawing down their fat and protein capital or by reducing allocation to future reproduction (Crémel, Festa-Bianchet, et al., 2026; Festa-Bianchet, 1998). We previously established that harsh snow conditions generate a persistent carry-over mass deficit reducing individual body mass the following autumn (Crémel, Festa-Bianchet, et al., 2026). Our path analyses provide new insights into this causal chain by showing that snow-induced mass loss also contributes to reduced female survival, although this pathway is cumulative and relatively weak. Previous studies on ungulates have argued that autumn body mass is a less reliable predictor of winter survival in adult males than in females, because heavier males invest more heavily in the rut (Mysterud et al., 2004; Preston et al., 2003). High reproductive effort can reduce or negate the survival benefits of a greater autumn mass (Forsyth et al., 2005). Variable mass loss during the rut (Pelletier & Festa-Bianchet, 2004) likely explains why indirect effects mediated by body mass was less certain in males. Although we detected a carry-over effect of previous snow conditions on survival via autumn body mass for both sexes, its small magnitude suggests that indirect winter pathways are far less influential than growing season conditions in driving mass-mediated survival in ungulates (Hurley et al., 2014). Our analyses revealed a direct effect of average snow depth that affects more vulnerable life stages, such as senescent individuals, more pronounced in males, the sex known to be more susceptible to resource scarcity (Clutton-Brock, Major, et al., 1985; Toïgo et al., 2007). A direct effect of snow depth on survival has similarly been documented in other ungulates (Horne et al., 2019; Jackson et al., 2021; LaSharr et al., 2023; Sirén et al., 2025).

The threshold analyses highlight how reliance on mean environmental conditions may fail to capture the acute demographic filters imposed by climate change. Deep snow triggers severe survival reductions across all individuals. Threshold effects could be linked to species-specific morphological limits. For instance, we found that survival strongly declines when snow depth exceeds 90 cm for adult females and lambs, compared to 100 cm for adult males. This threshold difference likely stems from sexual size dimorphism; the longer legs of rams (Blood et al., 1970) provide greater chest clearance (Telfer & Kelsall, 1984), allowing them to navigate deeper snow before reaching critical locomotor limitations. These critical depths correspond closely to the non-linear escalation of locomotory costs described by Dailey & Hobbs (1989). Given an average chest height of 40 to 50 cm for bighorn sheep, the metabolic cost of movement increases drastically up to an asymptote at approximately double of chest height. The 90-100 cm thresholds therefore correspond to the physical zone where locomotion costs quadruple (Dailey & Hobbs, 1989). Moreover, the high energetic cost of digging through snow (Goodson et al., 1991; Skogland, 1978) can prevent individuals from accessing forage under deep snow. These combined constraints likely drive the observed drops in survival in winters with deep snow (Jackson et al., 2021; Sirén et al., 2025). Furthermore, these physical limitations can induce compounding environmental pressures, including an increase in predation risk. Deep snow disproportionately impedes ungulate movement, potentially increasing their vulnerability to predators like wolves or cougars (Huggard, 1993; J drzejewski et al., 2002; Sullender et al., 2023). Because our models controlled for discrete, episodic predation events using a random year effect, the negative response reported here reflects a direct, snow-mediated constraint rather than isolated mortality pulses. Breaching morphological thresholds does not just create a direct energetic constraint but may also modify trophic interactions by intensifying prey vulnerability, a mechanism likely common to many snow ecosystems (Sullender et al., 2023).

As climate change broadly increases the frequency of extreme winter precipitation events (Lute et al., 2015; Wang et al., 2025), threshold effects become increasingly critical. Snowpack dynamics in the Rockies are highly spatially heterogeneous (Broberg, 2021; Lute et al., 2015; Wang et al., 2025); some sites are experiencing less snow, whereas others face heavier accumulations. At Ram Mountain, our multi-decadal reconstruction suggests a localized increasing trend in average snow depth (Crémel, Langlois, et al., 2026), primarily driven by the rise in deep snow events. Change in precipitation and climatic regime may alter the vital rates (Desforges et al., 2021) and threaten the long-term persistence of populations (White et al., 2025). Our analyses reveal that the increase in deep snow events over the last 45 years has occurred in parallel with a density-independent temporal decline in survival across all demographic groups. While lambs suffered the steepest demographic drop, adults were also profoundly impacted, experiencing a substantial erosion in survival that was particularly pronounced in females. This demographic erosion highlights the physiological limits of life-history buffering (Gaillard et al., 2000; Pfister, 1998). This pattern also illustrates a nonlinear demographic response increasingly recognized as central to buffering capacity (Gascoigne et al., 2025; Le Coeur et al., 2022). While selection favors allocation to somatic maintenance, this strategy offers diminishing protection when climate regimes shift beyond historical limits. The decline in adult female survival can precipitate rapid population declines due to its high elasticity on population growth (Gaillard et al., 2000). Ultimately, our results demonstrate that the widely accepted assumption of adult survival resilience may no longer hold under the increasing pressures of climate change.

## Conclusion

Our study demonstrates that predictions of population dynamics under climate change must account for non-linear physical thresholds. While adults of long-lived species can buffer average winter fluctuations through adaptations, this demographic resilience collapses when weather events breach critical morphological limits. As climate change intensifies the change in snow regime globally, previously buffered prime-aged cohorts may change, risking sudden population declines. Shifting attention from climatic means to frequency of events breaching species-specific morphological, energetic or physiological thresholds could provide deeper insights into wildlife population dynamics in an increasingly volatile climate.

## Acknowledgments

We are grateful to the many students, colleagues, and research assistants who have participated in data collection since 1971, with special thanks to J. Jorgenson. We also thank C. Feder and A. Hubbs for their continued support of the Ram Mountain research program.

## Author contributions

K.C., F.P., and M.F-B. conceptualized the study and performed data collection. K.C. performed the statistical analyses, interpreted the results, and drafted the manuscript. F.P., M.F-B., and A.L. contributed to the writing and critical revision of the manuscript. F.P. is responsible for the long-term research program at Ram Mountain.

## Data availability statement

Data and code are available in an Github repository https://github.com/Kallan-Cr/bighorn-survival-hierarchical

