## Appendix S1 for "Mortality in an alpine ungulate increases nonlinearly with snow depth"

**Table S1**: List of prediction tested for survival probability, along with the corresponding population group for bighorn sheep on Ram Mountain, Alberta, Canada

| Prediction | Population group |
| --- | --- |
| **1)** Survival probability was negatively associated with winter conditions. | Both |
| **2)** Survival probability was negatively associated with winter conditions through an interaction with age. | Males and females 1 year+ |
| **3)** Survival probability was negatively associated with winter conditions, with a stronger effect under high population density. | Both |

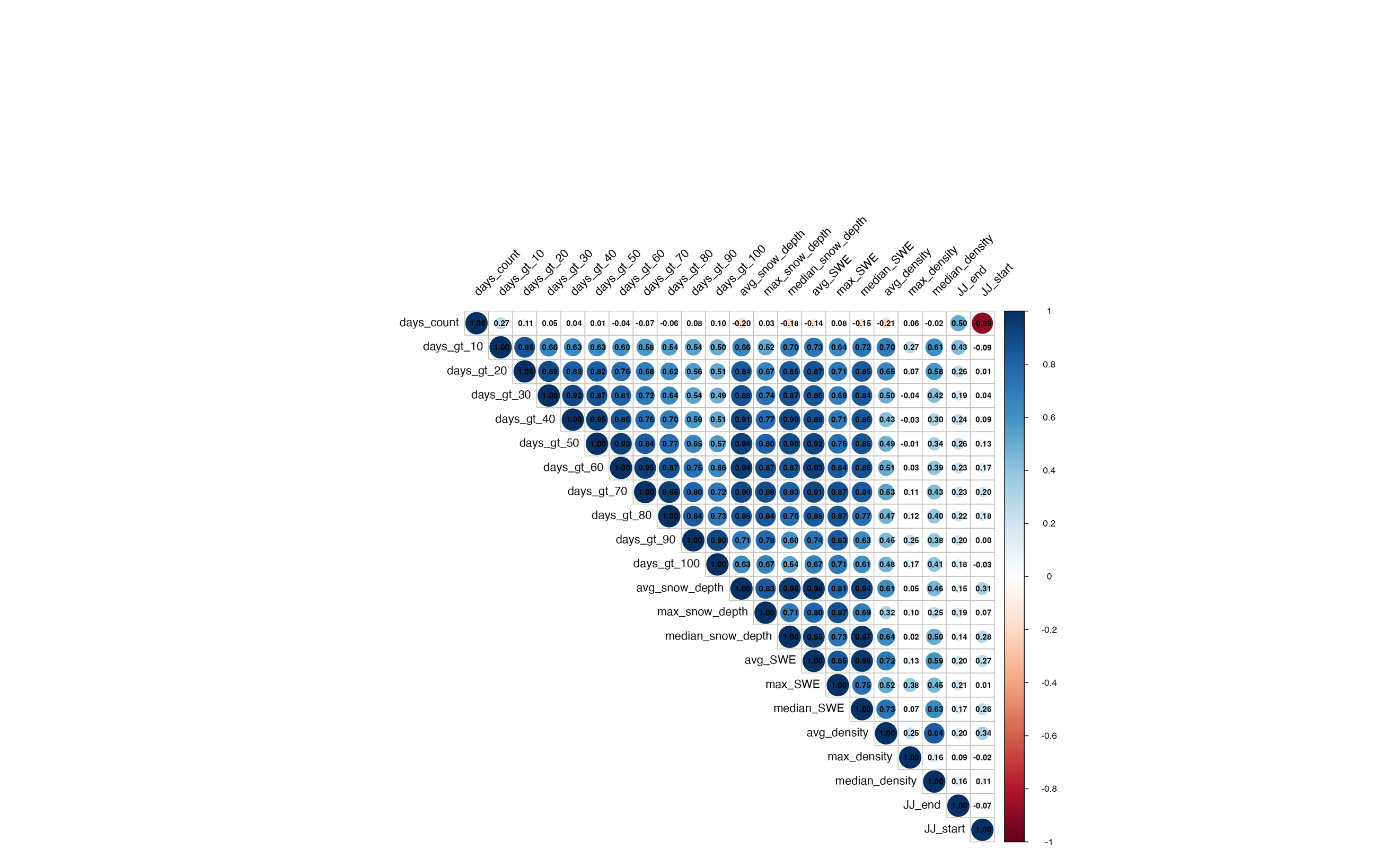

**Figure S1**: Correlation matrix between snow characteristic variables (duration of snow cover (days_count), snowdepth threshold (days_gt_*), mean, maximum, median of snow cover depth(*_snow_depth), mean, maximum, median of snow water equivalent (*_SWE) and mean, maximum and median of snow cover density (*_density)), end (JJ_end) and start date (JJ_start).

**Table S2**: Comparison of age modeling for females and males aged 1 year and over. For each population group, three types of modeling were tested: linear modeling, quadratic modeling and nonlinear modeling with a natural 3rd degree spline. Model comparison is based on ELPD. The model with the highest ELPD was selected as the best.

| **Population group** | **Model type** | **ELPD** | **∆ ELPD** |
| --- | --- | --- | --- |
| Female 1 yr+ | Linear | -714.5 | -9.1 |
| Female 1 yr+ | Quadratic | -708.8 | -3.4 |
| **Female 1 yr+** | **Spline 3^rd^ degree** | **-705.4** | **0.00** |
| Male 1 yr+ | Linear | -335.3 | -1.6 |
| Male 1 yr+ | Quadratic | -335.4 | -1.7 |
| **Male 1 yr+** | **Spline 3^rd^ degree** | **-333.8** | **0.00** |

**Table S3**: The modelling was performed separately for weaned lambs, adult females, and adult males. The table details the fixed effects structure within each model. Snow variables represent the set of three winter covariates: average snow depth, snow cover duration, and median snow density. All models included random intercepts for Year.

| **Groupe** | **Model** | **Fixed effect** |
| --- | --- | --- |
| Lambs | Base model | Survival ~ Sex * (Autumn mass + Population density + Snow variables) |
|  | Population density interaction with snow variables | Survival ~ Sex * Autumn mass + Sex * Population density * Snow variables |
| Females | Base model | Survival ~ Age (spline) + Autumn mass + Population density + Snow variables |
|  | Age interaction with snow variables | Survival ~ Population density + Autumn mass + Age (spline) * Snow variables |
|  | Population density interaction with snow variables | Survival ~ Age (spline) + Autumn mass + Population density * Snow variables |
| Males | Base model | Survival ~ Autumn mass + Age (spline) + Population density + Snow variables |
|  | Age interaction with snow variables | Survival ~ Population density + Autumn mass + Age (spline) * Snow variables |
|  | Population density interaction with snow variables | Survival ~ Autumn mass + Age (spline) + Population density * Snow variables |

**Table S4**: Comparison of models for each prediction (prediction number from Table S1) for each corresponding population group. Selection criteria include ELPD and ΔELPD. The model with the highest ELPD was selected as the best.

| **Population group** | **Prediction** | **ELPD** | **∆ELPD** |
| --- | --- | --- | --- |
| **Lamb** | **1** | -394.1 | **-0.0** |
| Lamb | 3 | -396.8 | -2.7 |
| **Female 1 yr+** | **1** | **-696.3** | **-0.0** |
| Female 1 yr+ | 2 | -701.5 | -5.2 |
| Female 1 yr+ | 3 | -697.6 | -1.3 |
| **Male 1 yr+** | **1** | **-336.8** | **-0.0** |
| Male 1 yr+ | 2 | -341.7 | -5.0 |
| Male 1 yr+ | 3 | -338.0 | -1.2 |

**Table S5**: Comparison of models for each snow depth threshold for each corresponding population group. Selection criteria include ELPD and ΔELPD. The model with the highest ELPD was selected as the best.

| **Population group** | **Prediction** | **ELPD** | **∆ELPD** |
| --- | --- | --- | --- |
| Lamb | **90** | -390.8 | 0.00 |
|  | 100 | -390.8 | -0.02 |
|  | 70 | -392.2 | -1.40 |
|  | 60 | -392.5 | -1.71 |
|  | 80 | -392.5 | -1.75 |
|  | 50 | -393.0 | -2.27 |
|  | 20 | -393.3 | -2.50 |
|  | 40 | -393.4 | -2.60 |
|  | 10 | -393.4 | -2.63 |
|  | 30 | -393.5 | -2.73 |
| Male | **100** | -335.7 | 0.00 |
|  | 50 | -335.8 | -0.15 |
|  | 90 | -335.8 | -0.18 |
|  | 60 | -336.1 | -0.43 |
|  | 40 | -336.1 | -0.45 |
|  | 30 | -336.1 | -0.46 |
|  | 20 | -336.2 | -0.52 |
|  | 10 | -336.2 | -0.53 |
|  | 70 | -336.2 | -0.54 |
|  | 80 | -336.2 | -0.58 |
| Female | **90** | -694.0 | 0.00 |
|  | 100 | -694.4 | -0.44 |
|  | 10 | -695.6 | -1.65 |
|  | 80 | -695.7 | -1.77 |
|  | 50 | -695.8 | -1.84 |
|  | 20 | -695.9 | -1.93 |
|  | 30 | -695.9 | -1.95 |
|  | 70 | -695.9 | -1.97 |
|  | 40 | -696.0 | -2.05 |
|  | 60 | -696.1 | -2.17 |
