## Appendix S2 for "Mortality in an alpine ungulate increases nonlinearly with snow depth"

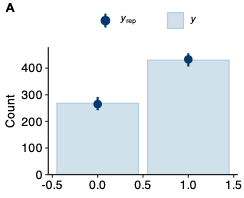

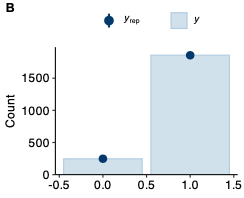

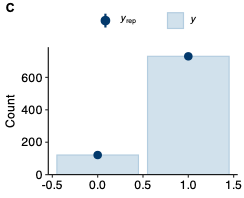

**Figure S1**: Posterior predictive check for the models assessing the effect of mean and median environmental variables on the survival of lambs (A), adult female (B) and adult male (C). The dark blue dots represent the observed counts in the data (y). The light blue bars represent the predicted counts from the 100 replicated datasets (yrep) drawn from the posterior predictive distribution. The close match between the observed data (dots) and the simulated replications (bars) indicates that the model adequately captures the distribution of the observed data.

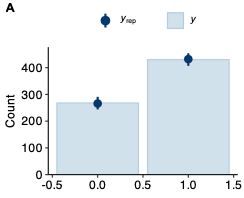

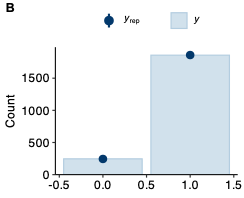

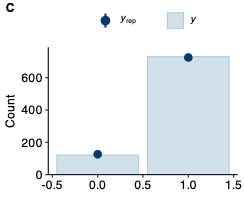

**Figure S2**: Posterior predictive check for the models assessing the effect of the snowdepth threshold on the survival of lambs (A), adult female (B) and adult male (C). The dark blue dots represent the observed counts in the data (y). The light blue bars represent the predicted counts from the 100 replicated datasets (yrep) drawn from the posterior predictive distribution. The close match between the observed data (dots) and the simulated replications (bars) indicates that the model adequately captures the distribution of the observed data.

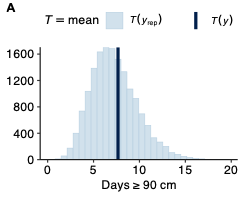

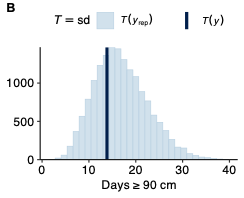

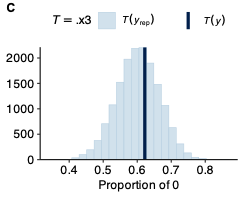

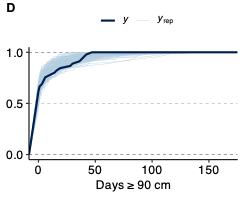

**Figure S3**: Posterior predictive checks for the beta-binomial model assessing the number of days exceeding a 90 cm threshold. Panels (A), (B), and (C) compare the observed test statistics (thick dark blue lines) to the expected distributions of the same statistics calculated from the replicated datasets (yrep, light blue bars). These panels represent the mean number of days, the standard deviation, and the proportion of zeros, respectively. Panel (D) displays the empirical cumulative distribution function (ECDF), where the thick dark blue line represents the observed data and the thin light blue lines represent the simulated replicated datasets drawn from the posterior predictive distribution. The strong alignment between the observed statistics and the simulated replication across all panels indicates that the model adequately captures the central tendency, dispersion, zero-inflation, and overall distribution of the observed data.

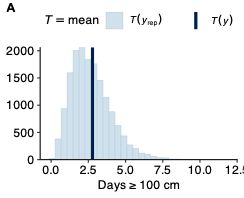

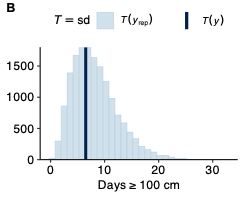

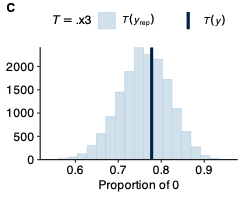

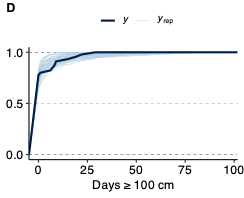

**Figure S4**: Posterior predictive checks for the beta-binomial model assessing the number of days exceeding a 100 cm threshold. Panels (A), (B), and (C) compare the observed test statistics (thick dark blue lines) to the expected distributions of the same statistics calculated from the replicated datasets (yrep, light blue bars). These panels represent the mean number of days, the standard deviation, and the proportion of zeros, respectively. Panel (D) displays the empirical cumulative distribution function (ECDF), where the thick dark blue line represents the observed data and the thin light blue lines represent the simulated replicated datasets drawn from the posterior predictive distribution. The strong alignment between the observed statistics and the simulated replication across all panels indicates that the model adequately captures the central tendency, dispersion, zero-inflation, and overall distribution of the observed data.

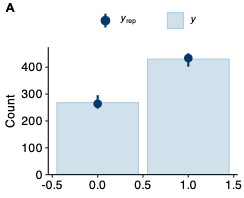

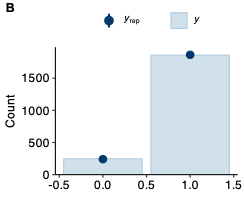

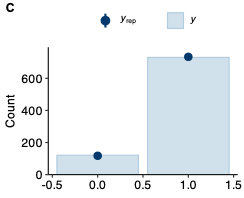

**Figure S5**: Posterior predictive check for the models assessing the temporal change in the survival of lambs (A), adult female (B) and adult male (C). The dark blue dots represent the observed counts in the data (y). The light blue bars represent the predicted counts from the 100 replicated datasets (yrep) drawn from the posterior predictive distribution. The close match between the observed data (dots) and the simulated replications (bars) indicates that the model adequately captures the distribution of the observed data.

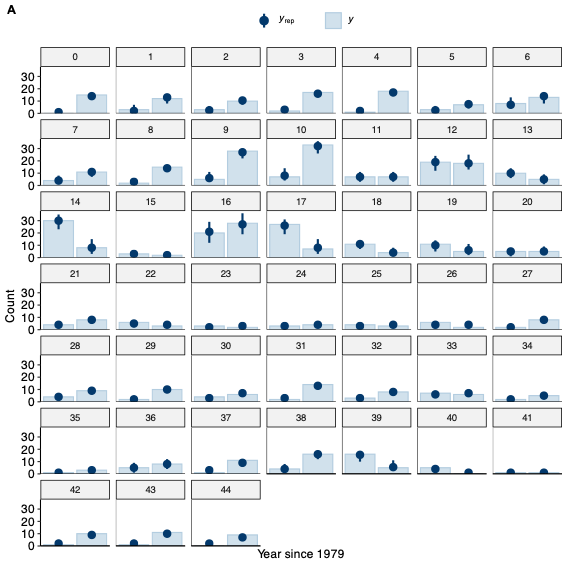

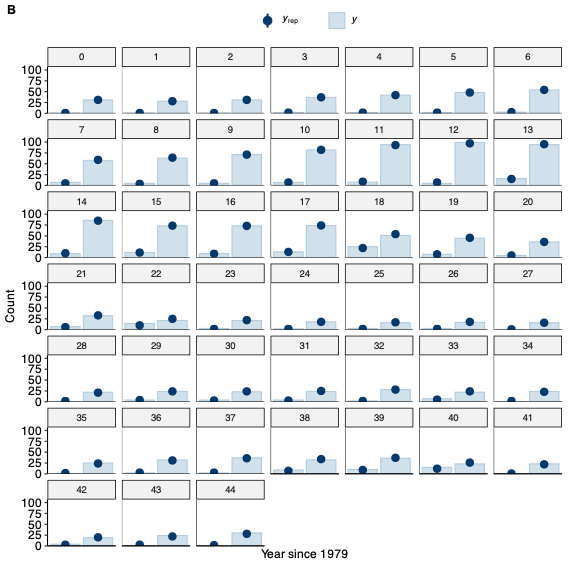

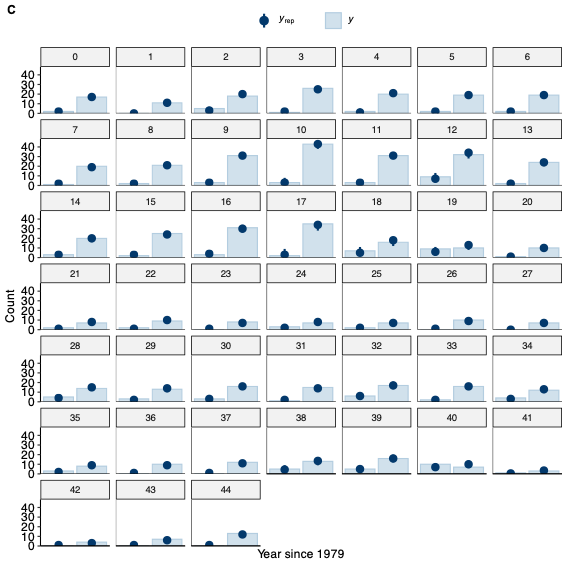

**Figure S6:** Posterior predictive checks for the temporal trend models assessing the survival of the lambs (A), adult females (B) and adult males (C). Each sub-panel represents a specific year of the study, indicated as the number of years since 1979. The dark blue dots represent the observed survival counts in the data (y). The light blue bars represent the predicted counts from the 100 replicated datasets (yrep) drawn from the posterior predictive distribution. The strong alignment between the observed data and the simulated replications across all years indicates that the models adequately capture the inter-annual fluctuations and temporal dynamics of survival for each group.
