## Appendix S3 for "Mortality in an alpine ungulate increases nonlinearly with snow depth"

**Table S1**: Posterior estimates (logit scale) from the Bayesian models for each sex-age group. For each fixed effect, posterior mean estimates, standard errors (Est.Error) and 95% credible intervals (l-95% CI to u-95% CI) are presented. Convergence diagnostics include the R-hat statistic (Rhat) and effective sample sizes for the bulk and tail of the posterior distributions (Bulk_ESS and Tail_ESS, respectively). All predictors were standardized prior to model fitting.

| **Population group** | **Fixed effect** | **Estimate** | **Est.Error** | **l-95% CI** | **u-95% CI** | **Rhat** | **Bulk_ESS** | **Tail_ESS** |
| --- | --- | --- | --- | --- | --- | --- | --- | --- |
| **Females** | Intercept | 3.566 | 0.389 | 2.813 | 4.335 | 1.0 | 13954 | 16500 |
|  | ns(Age) 1 | -2.249 | 0.490 | -3.206 | -1.293 | 1.0 | 23822 | 18558 |
|  | ns(Age) 2 | -4.087 | 0.949 | -5.956 | -2.229 | 1.0 | 15281 | 17224 |
|  | ns(Age) 3 | -3.567 | 0.642 | -4.841 | -2.290 | 1.0 | 20483 | 18688 |
|  | Autumn body mass | 0.793 | 0.164 | 0.472 | 1.116 | 1.0 | 15552 | 16742 |
|  | Population density | 0.008 | 0.149 | -0.295 | 0.296 | 1.0 | 10571 | 14648 |
|  | Avg. snow depth | -0.129 | 0.153 | -0.427 | 0.176 | 1.0 | 10647 | 14131 |
|  | Days count | -0.003 | 0.141 | -0.283 | 0.273 | 1.0 | 13277 | 16452 |
|  | Median density | -0.048 | 0.152 | -0.349 | 0.251 | 1.0 | 11506 | 15164 |
| **Males** | Intercept | 2.146 | 0.428 | 1.309 | 2.994 | 1.0 | 9546 | 13466 |
|  | Autumn body mass | 0.228 | 0.305 | -0.368 | 0.826 | 1.0 | 9222 | 13321 |
|  | Population density | 0.086 | 0.174 | -0.265 | 0.418 | 1.0 | 11764 | 15207 |
|  | Avg. snow depth | -0.183 | 0.190 | -0.556 | 0.188 | 1.0 | 11810 | 14139 |
|  | Days count | 0.113 | 0.170 | -0.223 | 0.446 | 1.0 | 13289 | 16014 |
|  | Median density | 0.061 | 0.190 | -0.313 | 0.431 | 1.0 | 12688 | 15570 |
|  | ns(Age) 1 | -1.353 | 1.089 | -3.501 | 0.783 | 1.0 | 12198 | 16578 |
|  | ns(Age) 2 | -1.057 | 1.264 | -3.526 | 1.442 | 1.0 | 8846 | 12051 |
|  | ns(Age) 3 | -2.133 | 1.135 | -4.352 | 0.086 | 1.0 | 12686 | 15899 |
| **Lambs** | Intercept | 0.945 | 0.222 | 0.516 | 1.393 | 1.0 | 10058 | 13423 |
|  | Sex (Male) | -0.790 | 0.204 | -1.193 | -0.394 | 1.0 | 25306 | 18917 |
|  | Avg. snow depth | -0.532 | 0.264 | -1.047 | -0.013 | 1.0 | 8866 | 12834 |
|  | Autumn body mass | 0.272 | 0.171 | -0.056 | 0.606 | 1.0 | 14601 | 16846 |
|  | Days count | -0.262 | 0.220 | -0.702 | 0.170 | 1.0 | 10134 | 13931 |
|  | Median density | 0.049 | 0.268 | -0.478 | 0.576 | 1.0 | 8552 | 12187 |
|  | Population density | -0.675 | 0.220 | -1.114 | -0.250 | 1.0 | 8393 | 12096 |
|  | Sex × Avg. snow depth | 0.307 | 0.250 | -0.178 | 0.801 | 1.0 | 12283 | 15488 |
|  | Sex × Autumn body mass | 0.359 | 0.217 | -0.071 | 0.788 | 1.0 | 15461 | 17035 |
|  | Sex × Days count | 0.163 | 0.204 | -0.237 | 0.564 | 1.0 | 17767 | 17848 |
|  | Sex × Median density | -0.294 | 0.251 | -0.789 | 0.196 | 1.0 | 12965 | 14887 |
|  | Sex × Population density | 0.416 | 0.198 | 0.028 | 0.803 | 1.0 | 18751 | 18163 |

**Table S2**: Posterior estimates (logit scale) from the Bayesian models for each sex-age group for the snow depth threshold analysis. For each fixed effect, posterior mean estimates, standard errors (Est.Error) and 95% credible intervals (l-95% CI to u-95% CI) are presented. Convergence diagnostics include the R-hat statistic (Rhat) and effective sample sizes for the bulk and tail of the posterior distributions (Bulk_ESS and Tail_ESS, respectively). All predictors were standardized prior to model fitting.

| **Population group** | **Fixed effect** | **Estimate** | **Est.Error** | **l-95% CI** | **u-95% CI** | **Rhat** | **Bulk_ESS** | **Tail_ESS** |
| --- | --- | --- | --- | --- | --- | --- | --- | --- |
| **Females** | Intercept | 3.569 | 0.374 | 2.849 | 4.315 | 1.0 | 14853 | 17422 |
|  | ns(Age) 1 | -2.266 | 0.490 | -3.226 | -1.303 | 1.0 | 26229 | 19185 |
|  | ns(Age) 2 | -4.126 | 0.933 | -5.970 | -2.329 | 1.0 | 15943 | 17400 |
|  | ns(Age) 3 | -3.604 | 0.646 | -4.877 | -2.348 | 1.0 | 22329 | 18249 |
|  | Autumn body mass | 0.789 | 0.162 | 0.475 | 1.108 | 1.0 | 16181 | 17356 |
|  | Population density | -0.055 | 0.128 | -0.313 | 0.190 | 1.0 | 13498 | 16626 |
|  | Median density | 0.084 | 0.132 | -0.178 | 0.341 | 1.0 | 16418 | 17768 |
|  | Days ≥ 90 cm | -0.381 | 0.115 | -0.605 | -0.154 | 1.0 | 15317 | 16633 |
| **Males** | Intercept | 2.177 | 0.422 | 1.353 | 3.032 | 1.0 | 8882 | 13160 |
|  | ns(Age) 1 | -1.495 | 1.082 | -3.604 | 0.627 | 1.0 | 11310 | 15689 |
|  | ns(Age) 2 | -1.137 | 1.244 | -3.559 | 1.311 | 1.0 | 8177 | 12743 |
|  | ns(Age) 3 | -2.074 | 1.106 | -4.237 | 0.112 | 1.0 | 12076 | 15772 |
|  | Autumn body mass | 0.253 | 0.303 | -0.339 | 0.849 | 1.0 | 8321 | 12445 |
|  | Population density | 0.049 | 0.171 | -0.298 | 0.376 | 1.0 | 12209 | 14267 |
|  | Median density | 0.120 | 0.188 | -0.252 | 0.488 | 1.0 | 12790 | 15274 |
|  | Days ≥ 100 cm | -0.281 | 0.170 | -0.626 | 0.046 | 1.0 | 11237 | 13772 |
| **Lambs** | Intercept | 0.959 | 0.216 | 0.543 | 1.397 | 1.0 | 10783 | 14898 |
|  | Sex (Male) | -0.793 | 0.202 | -1.194 | -0.400 | 1.0 | 30458 | 18974 |
|  | Autumn body mass | 0.312 | 0.171 | -0.024 | 0.646 | 1.0 | 15996 | 18069 |
|  | Median density | 0.124 | 0.248 | -0.368 | 0.606 | 1.0 | 9551 | 13696 |
|  | Population density | -0.752 | 0.214 | -1.182 | -0.340 | 1.0 | 9662 | 14229 |
|  | Days ≥ 90 cm | -0.659 | 0.225 | -1.112 | -0.221 | 1.0 | 9712 | 13704 |
|  | Sex × Autumn body mass | 0.329 | 0.215 | -0.092 | 0.747 | 1.0 | 17539 | 18893 |
|  | Sex × Median density | -0.381 | 0.235 | -0.842 | 0.079 | 1.0 | 15098 | 17830 |
|  | Sex × Population density | 0.507 | 0.201 | 0.119 | 0.908 | 1.0 | 18143 | 17995 |
|  | Sex × Days ≥ 90 cm | 0.475 | 0.221 | 0.047 | 0.913 | 1.0 | 16410 | 17902 |

**Table S3**: Posterior estimates from the Bayesian models for each selected snow depth threshold. For each fixed effect, posterior mean estimates, standard errors (Est.Error) and 95% credible intervals (l-95% CI to u-95% CI) are presented. Convergence diagnostics include the R-hat statistic (Rhat) and effective sample sizes for the bulk and tail of the posterior distributions (Bulk_ESS and Tail_ESS, respectively).

| **Threshold** | **Fixed effect** | **Estimate** | **Est.Error** | **l-95% CI** | **u-95% CI** | **Rhat** | **Bulk_ESS** | **Tail_ESS** |
| --- | --- | --- | --- | --- | --- | --- | --- | --- |
| **Days ≥ 90 cm** | Intercept | -5.961 | 0.117 | -6.192 | -5.734 | 1.0 | 6623 | 8410 |
|  | Year | 0.076 | 0.003 | 0.069 | 0.082 | 1.0 | 8922 | 10009 |
| **Days ≥ 100 cm** | Intercept | -6.867 | 0.157 | -7.176 | -6.563 | 1.0 | 6469 | 7622 |
|  | Year | 0.072 | 0.004 | 0.064 | 0.080 | 1.0 | 8325 | 8872 |

**Table S4**: Posterior estimates (logit scale) from the Bayesian models for each sex-age group for the survival trend analysis. For each fixed effect, posterior mean estimates, standard errors (Est.Error) and 95% credible intervals (l-95% CI to u-95% CI) are presented. Convergence diagnostics include the R-hat statistic (Rhat) and effective sample sizes for the bulk and tail of the posterior distributions (Bulk_ESS and Tail_ESS, respectively). All predictors were standardized prior to model fitting.

| **Population group** | **Fixed effect** | **Estimate** | **Est.Error** | **l-95% CI** | **u-95% CI** | **Rhat** | **Bulk_ESS** | **Tail_ESS** |
| --- | --- | --- | --- | --- | --- | --- | --- | --- |
| **Females** | Intercept | 4.102 | 0.453 | 3.231 | 4.999 | 1.0 | 16858 | 18168 |
|  | ns(Age) 1 | -2.218 | 0.486 | -3.176 | -1.269 | 1.0 | 27538 | 19030 |
|  | ns(Age) 2 | -4.066 | 0.942 | -5.909 | -2.232 | 1.0 | 18106 | 17591 |
|  | ns(Age) 3 | -3.545 | 0.639 | -4.808 | -2.286 | 1.0 | 24476 | 19419 |
|  | Autumn body mass | 0.790 | 0.162 | 0.474 | 1.112 | 1.0 | 17096 | 17902 |
|  | Population density | -0.169 | 0.155 | -0.480 | 0.130 | 1.0 | 13232 | 17275 |
|  | Year | -0.030 | 0.012 | -0.054 | -0.005 | 1.0 | 17288 | 18286 |
| **Males** | Intercept | 2.421 | 0.469 | 1.516 | 3.340 | 1.0 | 12388 | 16216 |
|  | ns(Age) 1 | -1.171 | 1.084 | -3.288 | 0.967 | 1.0 | 12499 | 15985 |
|  | ns(Age) 2 | -0.897 | 1.235 | -3.246 | 1.525 | 1.0 | 9265 | 14173 |
|  | ns(Age) 3 | -2.033 | 1.100 | -4.145 | 0.162 | 1.0 | 13623 | 17425 |
|  | Autumn body mass | 0.190 | 0.298 | -0.393 | 0.767 | 1.0 | 9545 | 14117 |
|  | Population density | -0.015 | 0.192 | -0.401 | 0.358 | 1.0 | 10860 | 13765 |
|  | Year | -0.018 | 0.015 | -0.047 | 0.011 | 1.0 | 12296 | 15532 |
| **Lambs** | Intercept | 1.739 | 0.484 | 0.815 | 2.718 | 1.0 | 8775 | 13426 |
|  | Sex (Male) | -1.036 | 0.464 | -1.980 | -0.154 | 1.0 | 10061 | 15161 |
|  | Autumn body mass | 0.290 | 0.168 | -0.037 | 0.625 | 1.0 | 15865 | 17844 |
|  | Population density | -0.906 | 0.256 | -1.413 | -0.414 | 1.0 | 7281 | 12120 |
|  | Year | -0.041 | 0.022 | -0.084 | 0.000 | 1.0 | 8398 | 13130 |
|  | Sex × Autumn body mass | 0.318 | 0.214 | -0.097 | 0.743 | 1.0 | 17036 | 17850 |
|  | Sex × Population density | 0.466 | 0.235 | 0.015 | 0.930 | 1.0 | 11823 | 16848 |
|  | Sex × Year | 0.013 | 0.021 | -0.028 | 0.055 | 1.0 | 9771 | 14287 |

**Table S5**: G-computation results summarizing the estimated causal effects of each snow variable on survival probability, by sex and age. For each snow variable, population group, and scenario, the slope estimates and its 95% credible interval (l-95% CI, u-95% CI) are shown alongside the probability of direction (Pd) and the absolute difference in survival.

| **Snow variables** | **Sex** | **Age-classes** | **Scenarios** | **Slopes** | **l-95% CI** | **u-95% CI** | **Pd** | **Absolute differences** |
| --- | --- | --- | --- | --- | --- | --- | --- | --- |
| **Days count** | **Female** | Yearling | Direct effect | 0.0000 | −0.0009 | 0.0014 | 0.51 | 0.0000 |
|  |  |  | Indirect effect (mediation) | −0.0000 | −0.0008 | −0.0000 | 0.99 | −0.0033 |
|  |  |  | Total effect | −0.0000 | −0.0013 | 0.0009 | 0.69 | −0.0007 |
|  |  | Prime | Direct effect | 0.0000 | −0.0009 | 0.0015 | 0.51 | 0.0000 |
|  |  |  | Indirect effect (mediation) | −0.0000 | −0.0004 | 0.0000 | 0.90 | −0.0019 |
|  |  |  | Total effect | −0.0000 | −0.0011 | 0.0013 | 0.58 | −0.0001 |
|  |  | Senescent | Direct effect | 0.0000 | −0.0011 | 0.0015 | 0.51 | 0.0000 |
|  |  |  | Indirect effect (mediation) | −0.0000 | −0.0004 | 0.0000 | 0.91 | −0.0011 |
|  |  |  | Total effect | −0.0000 | −0.0012 | 0.0012 | 0.58 | −0.0001 |
|  | **Male** | Yearling | Direct effect | 0.0000 | −0.0005 | 0.0025 | 0.78 | 0.0028 |
|  |  |  | Indirect effect (mediation) | −0.0000 | −0.0002 | 0.0000 | 0.79 | −0.0002 |
|  |  |  | Total effect | 0.0000 | −0.0005 | 0.0024 | 0.75 | 0.0024 |
|  |  | Prime | Direct effect | 0.0000 | −0.0004 | 0.0025 | 0.78 | 0.0026 |
|  |  |  | Indirect effect (mediation) | −0.0000 | −0.0002 | 0.0000 | 0.78 | −0.0001 |
|  |  |  | Total effect | 0.0000 | −0.0006 | 0.0024 | 0.75 | 0.0022 |
|  |  | Senescent | Direct effect | 0.0009 | −0.0019 | 0.0036 | 0.78 | 0.1052 |
|  |  |  | Indirect effect (mediation) | −0.0001 | −0.0005 | 0.0001 | 0.79 | −0.0100 |
|  |  |  | Total effect | 0.0008 | −0.0021 | 0.0034 | 0.75 | 0.0921 |
| **Snow depth** | **Female** | Yearling | Direct effect | −0.0012 | −0.0044 | 0.0013 | 0.82 | −0.0524 |
|  |  |  | Indirect effect (mediation) | −0.0005 | −0.0014 | −0.0001 | 0.99 | −0.0236 |
|  |  |  | Total effect | −0.0020 | −0.0056 | 0.0007 | 0.92 | −0.0897 |
|  |  | Prime | Direct effect | −0.0011 | −0.0041 | 0.0012 | 0.82 | −0.0454 |
|  |  |  | Indirect effect (mediation) | −0.0002 | −0.0007 | 0.0001 | 0.95 | −0.0094 |
|  |  |  | Total effect | −0.0014 | −0.0046 | 0.0010 | 0.87 | −0.0603 |
|  |  | Senescent | Direct effect | −0.0020 | −0.0066 | 0.0021 | 0.82 | −0.0891 |
|  |  |  | Indirect effect (mediation) | −0.0003 | −0.0009 | 0.0002 | 0.86 | −0.0125 |
|  |  |  | Total effect | −0.0024 | −0.0071 | 0.0019 | 0.86 | −0.1053 |
|  | **Male** | Yearling | Direct effect | −0.0018 | −0.0070 | 0.0017 | 0.82 | −0.0819 |
|  |  |  | Indirect effect (mediation) | −0.0001 | −0.0007 | 0.0002 | 0.80 | −0.0056 |
|  |  |  | Total effect | −0.0021 | −0.0075 | 0.0017 | 0.84 | −0.0927 |
|  |  | Prime | Direct effect | −0.0013 | −0.0054 | 0.0012 | 0.82 | −0.0565 |
|  |  |  | Indirect effect (mediation) | −0.0002 | −0.0008 | 0.0003 | 0.80 | −0.0075 |
|  |  |  | Total effect | −0.0016 | −0.0058 | 0.0012 | 0.86 | −0.0703 |
|  |  | Senescent | Direct effect | −0.0027 | −0.0097 | 0.0026 | 0.82 | −0.1211 |
|  |  |  | Indirect effect (mediation) | −0.0004 | −0.0020 | 0.0004 | 0.80 | −0.0181 |
|  |  |  | Total effect | −0.0033 | −0.0103 | 0.0021 | 0.86 | −0.1447 |
| **Median density** | **Male** | Prime | Direct effect | 0.0000 | −0.0016 | 0.0014 | 0.64 | 0.0001 |
|  |  |  | Indirect effect (mediation) | 0.0000 | −0.0001 | 0.0002 | 0.75 | 0.0001 |
|  |  |  | Total effect | 0.0000 | −0.0015 | 0.0014 | 0.65 | 0.0002 |
|  |  | Senescent | Direct effect | 0.0005 | −0.0028 | 0.0042 | 0.64 | 0.0361 |
|  |  |  | Indirect effect (mediation) | 0.0000 | −0.0002 | 0.0005 | 0.70 | 0.0025 |
|  |  |  | Total effect | 0.0005 | −0.0027 | 0.0042 | 0.65 | 0.0390 |
